# A Novel Non-hypermucoviscous ST11 Hypervirulent Carbapenem-resistant *Klebsiella pneumoniae* Lacking Classical Virulence Factors

**DOI:** 10.1101/2020.02.28.970863

**Authors:** Min Zhang, Jinyong Zhang, Jia Li, Shuye Xu, Chunxia Hu, Qiuyang Deng, Xianglin Wu, Lihua Xiao, Zhou Liu, Xiyao Yang, Xiaoqian Hu, Wenhui Zhang, Nan Wang, Liqi Yang, Shiyi Liu, Ruiqin Cui, Zhen Hui, Yuxin Zhong, Yutian Luo, Huaisheng Chen, Weiyuan Wu, Jinsong Wu, Yuemei Lu, Xueyan Liu, Quanming Zou, Wei Huang

## Abstract

**Background:** Hypervirulent *Klebsiella pneumoniae* lacking classical virulence factors is uncommon, and the virulence mechanisms of this organism are not understood.

**Methods:** Following a retrospective study of carbapenem-resistant *K. pneumoniae* based on core genome multilocus sequence typing (cgMLST), isolates that caused high mortality were investigated with a genome-wide association study (GWAS), proteome analysis and an animal model.

**Results:** The sublineage of sequence type 11 (ST11) *K. pneumoniae*, which belongs to complex type 3176 (CT3176) and K-locus 47 (KL47), was highlighted due to the high mortality of infected patients. GWAS analysis showed that *ampR* was associated with the CT3176 isolates. In a mouse model, the mortality of *ampR*-carrying isolates was comparable to that of the typical hypervirulent isolate GM2. Even during the first 24 hours of infection, the bacterial load and pathological changes of the *ampR*-carrying isolates in the lungs were more severe than those of GM2. The *ampR* complement mutant was able to enhance the virulence of the KL47 isolate but not the virulence of KL1. Proteome analysis showed that the expression of WcaJ in the *ampR*^+^ isolates was significantly higher than that in the *ampR*^−^ isolates, and this result was also confirmed by transcription tests and capsule staining. It is suggested that the enhancement of the initial stage of capsule synthesis may be the cause of the high virulence of these non-hypermucoviscous ST11 carbapenem-resistant *K. pneumoniae* isolates.

**Conclusions:** Non-hypermucoviscous ST11 hypervirulent carbapenem-resistant *K. pneumoniae* warrants continued surveillance and investigation.

---

Classical *K. pneumoniae* (cKp) and hypervirulent *K. pneumoniae* (hvKp) are two global *K. pneumoniae* pathotypes presently circulating [1, 2]. In North America and Europe, most *K. pneumoniae* infections are caused by cKp isolates, which are most commonly opportunistic pathogens causing infections primarily in the health care setting in hosts with comorbidities, who are immunocompromised, or who have existing wounds. The most problematic characteristic of this pathotype is the ability to acquire an increasing number of elements that confer antimicrobial resistance. Carbapenem-resistant *K. pneumoniae* (CRKP) associated with plasmid-encoded carbapenemases poses distinct clinical challenges and produces invasive infections with high mortality [3, 4].

Reports from Taiwan described a unique clinical syndrome of community-acquired, tissue-invasive *K. pneumoniae* infection in healthy individuals that often presented at multiple sites or subsequently spread, including pyogenic liver abscesses in the absence of biliary tract disease, abscesses at non-hepatic sites, pneumonia, endophthalmitis, meningitis, and necrotizing fasciitis [5, 6]. hvKp was designated to distinguish this pathotype from cKp, and the incidence of infections due to hvKp has been steadily increasing over the last 3 decades in Asian Pacific countries [7–11]. Usually, most hvKp isolates are susceptible to antimicrobials [12]. However, the hvKp isolate has the potential to acquire drug resistance genes that encode extended-spectrum β-lactamases and carbapenemases by plasmid transformation, which was accomplished experimentally [13]. Antimicrobial-resistant hvKp was also observed in the clinical setting [14]. Furthermore, an extensively drug-resistant (XDR) cKp isolate that acquired part of an hvKp virulence plasmid caused a fatal nosocomial outbreak [15].

A characteristic that was initially believed to be sensitive and specific for hvKp isolates was a hypermucoviscous phenotype, which was defined by a positive string test [16]. When this phenotype was used alone to define an hvKp isolate, the misperception that occurred created some confusion in the literature as not all hvKp isolates were determined as being hypermucoviscous, and some cKp isolates possessed this characteristic [17]. Recently, multiple biomarkers including *peg-344*, *iroB*, *iucA*, plasmid-encoded *rmpA* and *rmpA2* and quantitative siderophore production have been shown to accurately predict hvKp isolates; these biomarkers could be used to develop a diagnostic test for use by clinical laboratories for optimal patient care and for use in epidemiologic surveillance and research [18].

Whether a hvKP or cKP isolate has hypervirulence due to acquired virulence plasmids, the presence of classical virulence factors is common. However, in this study, we identified a novel group of non-hypermucoviscous hypervirulent CRKP isolates lacking most classical virulence genes. A pan genome-wide association study (Pan-GWAS), proteome analysis and virulence test in an animal model revealed that AmpR improves the virulence of these isolates by regulating the initiation of capsule synthesis.

## Methods

### Clinical Isolates and Data Collection

Non-repetitive clinical *K. pneumoniae* isolates were collected from 5 hospitals in Guangdong and Anhui provinces between 2018 and 2019. All isolates were stocked at −80 °C prior to use. Clinical information on patients was obtained. Species identification and antimicrobial susceptibility testing were performed with the VITEK-2 compact system (bioMérieux, Marcy-l’Étoile, France). The results were interpreted in accordance with guidelines published by the Clinical and Laboratory Standards Institute (CLSI; document M100-S26) [19]. The identified species of all isolates were confirmed with matrix-assisted laser desorption/ionization mass spectrometry (bioMérieux, Marcy-l’Étoile, France). CRKP was defined as resistant to imipenem or meropenem.

### Hypermucoviscous Phenotypic Characterization

To identify the hypermucoviscous phenotype with the string test, isolates were inoculated onto agar plates containing 5% sheep blood and incubated at 37 °C overnight. The string test was deemed positive when a viscous string longer than 5 mm could be generated by touching a single colony with a standard inoculation loop and pulling the colony upwards [2].

### Whole Genome Sequencing (WGS)

A 1 mL culture volume (optical density at 600 nm [OD_600_] of 0.6) was used for genomic DNA (gDNA) extraction (Sangon, Shanghai, China). gDNA was sequenced on an Illumina HiSeq 2500 sequencer (Illumina, San Diego, CA, USA) using paired-end 150-bp reads. Genome assembly was performed using de novo SPAdes Genome Assembler (version 3.12.0) [20]. We performed capsule typing on the assembled sequences with Kaptive (version 0.5.1) [21]. Antimicrobial resistance genes and virulence factors were identified in the isolates by scanning the genome contigs against the ResFinder and VFDB databases using ABRicate (version 0.8.7). Multilocus sequence typing (MLST) and core genome MLST (cgMLST) genotyping analysis were performed with Ridom SeqSphere^+^ (version 5.1.0) [22]. Complex type (CT) was defined by following the *K. pneumoniae* cgMLST scheme (www.cgmlst.org/ncs/schema/2187931). Phylogenetic tree construction based on core genome single nucleotide variants (SNVs) was performed using the Harvest suite (version 1.2) with a 1,000-bootstrap test [23]. The online tool iTOL was used to display, manipulate and annotate the phylogenetic tree [24]. We annotated the genome sequences with Prokka (version 1.13.3) [25].

### Pan Genome-wide association study (Pan-GWAS) Analysis

We used the pan genome pipeline Roary (version 3.12.0) to put annotated assemblies in GFF3 format (produced by Prokka) and calculated the pan genome [26]. We used Scoary (version 1.6.16) to take the file from Roary, created a trait file and calculated the associations between all genes in the accessory genome and the traits. We reported a list of genes sorted by strength of association with *P*-value < 0.01 adjusted with Benjamini-Hochberg’s method for multiple comparisons correction [27].

### Construction of The *ampR* Complement Mutant

*K. pneumoniae ampR* (Supplementary Materials) was synthesized and flanked with 5’XbaI and 3’HindIII restriction sites. This fragment was restriction digested and cloned into pUC57. Recombinant DNA was recovered in *Escherichia coli* DH5α with ampicillin selection. After restriction digestion, the *ampR* fragment was ligated to the pBAD33 expression vector (Invitrogen) according to the manufacturer’s instructions. pBAD33-*ampR* was used for transformation using electroporation as a standard for laboratory-isolated *E. coli*. The *ampR* complement strains were confirmed by polymerase chain reaction (PCR) (Supplementary Table 1). The expression of *ampR* was induced by adding 100 μg/mL arabinose.

### Mouse Pulmonary Infection Models

A pneumonia model of *K. pneumoniae* in mice was used to test the virulence of isolates. Six- to eight-week-old female BALB/c mice under specific pathogen-free grade were purchased from Hunan SJA Laboratory Animal Co., Ltd. The mice were intraperitoneally anaesthetized with pentobarbital sodium (75 mg/kg) and inoculated with 1.0 × 10^7^ colony-forming units (CFU) of *K. pneumoniae* by non-invasive intratracheal instillation under direct vision. The survival of the mice was observed for 7 days post-infection. To assess bacterial burden in the lung, the organs from infected mice were homogenized and dissolved in 1 mL of phosphate buffer solution (PBS); 50 μL was inoculated onto liquid broth (LB) agar plates, and CFU enumeration was performed. For histopathology analysis, segments of the lungs were fixed with 10% neutral formalin, embedded in paraffin and stained with haematoxylin and eosin for visualization by light microscopy. Bacterial burden and histopathology analysis were performed after 24 hours of infection.

All animal care and use protocols in this study were performed in accordance with the Regulations for the Administration of Affairs Concerning Experimental Animals approved by the State Council of the People’s Republic of China. All animal experiments in this study were approved by the Animal Ethical and Experimental Committee of the Army Military Medical University (Chongqing, Permit No. 2011-04) in accordance with their rules and regulations.

### Proteome Analysis

A 1 mL culture volume (optical density at 600 nm [OD600] of 0.6) was used for protein extraction. The sample was sonicated three times on ice using a high-intensity ultrasonic processor (Scientz) in lysis buffer (8 M urea, 1% protease inhibitor cocktail). The remaining debris was removed by centrifugation at 12,000 g at 4 °C for 10 min. Finally, the supernatant was collected, and the protein concentration was determined with a BCA kit according to the manufacturer’s instructions. For digestion, the protein solution was reduced with 5 mM dithiothreitol for 30 min at 56 °C and alkylated with 11 mM iodoacetamide for 15 min at room temperature in darkness. The protein sample was then diluted by adding 100 mM TEAB with urea at a concentration of less than 2 M. Finally, trypsin was added at a 1:50 trypsin-to-protein mass ratio for the first digestion overnight and a 1:100 trypsin-to-protein mass ratio for a second 4 hours digestion. The tryptic peptides were dissolved in 0.1% formic acid (solvent A) and directly loaded onto a homemade reversed-phase analytical column (15-cm length, 75 μm i.d.). The gradient was performed as follows: an increase from 6% to 23% solvent B (0.1% formic acid in 98% acetonitrile) over 26 min, an increase from 23% to 35% solvent B in 8 min, an increase to 80% solvent B in 3 min and holding at 80% solvent B for the last 3 min at a constant flow rate of 400 nL/min on an EASY-nLC 1000 UPLC system. The peptides were subjected to NSI source followed by tandem mass spectrometry (MS/MS) in Q ExactiveTM Plus (Thermo) coupled online to the UPLC. The electrospray voltage applied was 2.0 kV. The m/z scan range was 350 to 1800 for full scan, and intact peptides were detected in the Orbitrap at a resolution of 70,000. Peptides were then selected for MS/MS using the NCE setting of 28, and the fragments were detected in the Orbitrap at a resolution of 17,500. The data-dependent procedure alternated between one MS scan followed by 20 MS/MS scans with 15.0 s dynamic exclusion. Automatic gain control (AGC) was set at 5E4. The fixed first mass was set as 100 m/z. The resulting MS/MS data were processed using the MaxQuant search engine (v.1.5.2.8). We used InterProScan (version 5.37-76.0) to annotate protein domains. A corrected *p*-value < 0.05 and a fold change > 1.2 were considered significant.

### Real-time quantitative PCR (RT-qPCR)

A 1 mL culture volume (optical density at 600 nm [OD_600_] of 0.6) was used for total RNA extraction (Sangon, Shanghai, China). cDNA was generated from 500 ng of total RNA. SYBR Premix Ex Taq II (Sangon, Shanghai, China) was used for qPCR. The primers used are listed in Supplementary Table 1. The 16S rRNA was used as a reference gene for normalization of expression levels.

### Capsule Staining

Bacterial capsules were detected using TyLer methods according to the manufacturer’s instructions (Solarbio, Beijing, China). Briefly, the bacterial smear was stained with crystal violet dye for 5-7 min and then rinsed with copper sulfate 2-3 times. The bacteria were observed and photographed under light microscopy (1000 ×).

### Statistical Analysis

Statistical analysis was performed using GraphPad Prism software version 8 (La Jolla, California).

### Data Availability

We deposited the genome sequences in GenBank under BioProject PRJNA517992. The genome sequence of GM2 was deposited in GenBank under BioProject PRJNA556307.

## Results

### Isolate Characteristics

Non-repetitive CRKP (n = 135) isolates were confirmed in this study. All isolates were assigned to sequence type 11 (ST11) and carried *K. pneumoniae* carbapenemase (KPC-2), and 1 isolate carried both OXA-23 and KPC-2 (Supplementary Figure 1). Isolates were assigned to 16 CTs using the cgMLST scheme. The most common CTs were CT1313 (n = 35), CT1291 (n = 20), CT3176 (n = 14), CT1814 (n = 13), and CT2410 (n = 11), followed by 11 other CTs identified in less than 10 isolates each (Figure 1). All isolates were assigned to two K-locus types, KL64 (n = 86) and KL47 (n = 49) (Supplementary Figure 2). No hypermucoviscous isolates were found. In the virulence gene analysis, we found that no isolates carrying multiple biomarkers for differentiation of the hvKp and cKp isolates had been reported previously (Figure 1, Supplementary Figure 2) [18].

**Figure 1.**
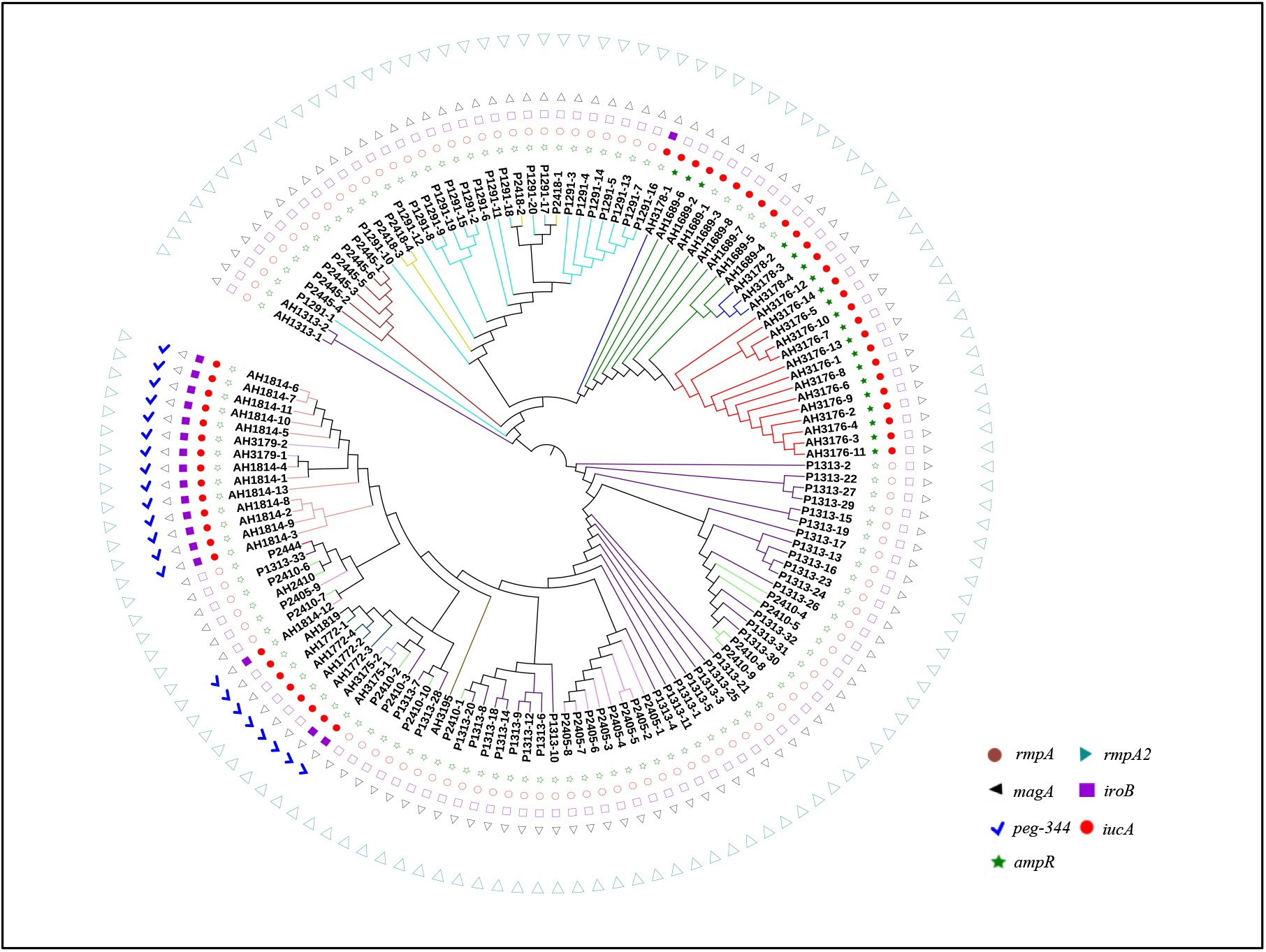
Phylogenetic analysis of 135 isolates in this study. The branches of different CT type isolates are shown in colours. The corresponding isolates of each CT type are CT1291 (P1291-1 to P1291-20), CT1313 (P1313-1 to P1313-33, AH1313-1 to AH1313-2), CT1689 (AH1689-1 to AH1689-8), CT1772 (AH1772-1 to AH1772-4), CT1814 (AH1814-1 to AH1814-13), CT1819 (AH1819), CT2405 (P2405-1 to P2405-9), CT2410 (P2410-1 to P2410-11), CT2418 (P2418-1 to P2418-4), CT2444 (P2444), CT2445 (P2445-1 to P2445-6), CT3175 (AH3175-1 to AH3175-2), CT3176 (AH3176-1 to AH3176-14), CT3178 (AH3178-1 to AH3178-4), CT3179 (AH3179-1 to AH3179-2), and CT3195 (AH3195). *rmpA*, *rmpA2*, *magA iroB*, *peg-344* and *iucA* were identified for the differentiation of hypervirulent *K. pneumoniae* from classical *K. pneumoniae* in a previous study [18]. Abbreviations: *rmpA/rmpA2*, cps transcriptional activator; *magA*, outer membrane protein; *iroB*, iron acquisition system; *peg-344*, metabolite transporter; *iucA*, aerobactin-related genes.

### Clinical Characteristics

There were no significant differences between the CTs in terms of demographics, infection types, main comorbidities, invasive operations, Charlson comorbidity index and Pitt bacteraemia score, except for central venous catheter, and all cause in-hospital mortality (*P* = 0.035 and *P* = 0.001, Fisher’s exact test). Patients infected with CT3176 had significantly higher mortality rates (78.6% vs 37.1%, *P* = 0.012; 78.6% vs 20.0%, *P* = 0.001; 78.6% vs 7.7%, *P* = 0.0003; 78.6% vs 31.0%, *P* = 0.004; Fisher’s exact test) compared with the CT1313, CT1291, CT1814 and the other CT groups, respectively. The utilization of central venous catheters in CT1814 was significantly higher than that in CT1313 and CT1291 (84.6% vs 51.4%, *P* = 0.049; 84.6% vs 35.0%, *P* = 0.011, Fisher’s exact test) (Table 1).

**Table 1.**
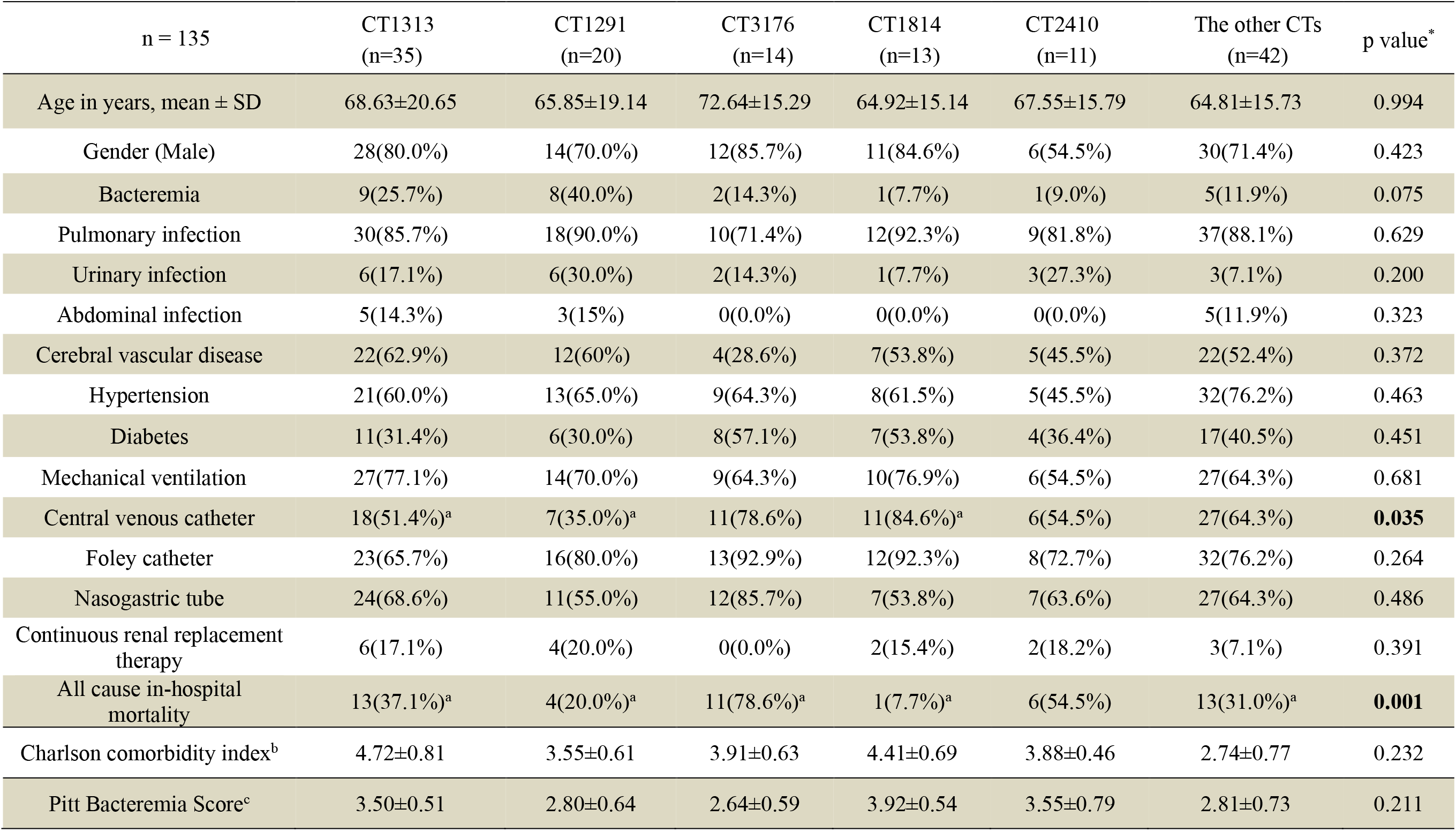
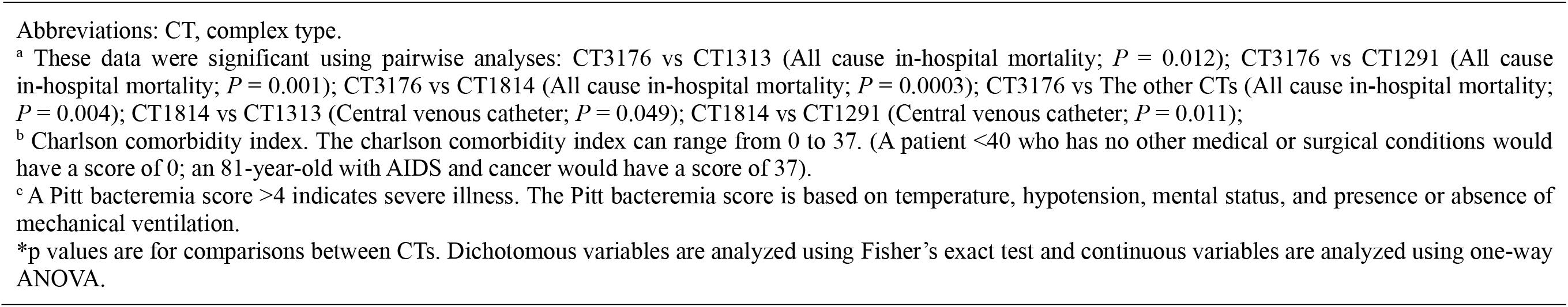
Demographics, comorbidities, invasive procedures and mortality of 135 patients with carbapenem-resistant *Klebsiella pneumoniae* infections in this study.

| n = 135 | CT1313<br>(n=35) | CT1291<br>(n=20) | CT3176<br>(n=14) | CT1814<br>(n=13) | CT2410<br>(n=11) | The other CTs<br>(n=42) | p value* |
| --- | --- | --- | --- | --- | --- | --- | --- |
| Age in years, mean ± SD | 68.63±20.65 | 65.85±19.14 | 72.64±15.29 | 64.92±15.14 | 67.55±15.79 | 64.81±15.73 | 0.994 |
| Gender (Male) | 28(80.0%) | 14(70.0%) | 12(85.7%) | 11(84.6%) | 6(54.5%) | 30(71.4%) | 0.423 |
| Bacteremia | 9(25.7%) | 8(40.0%) | 2(14.3%) | 1(7.7%) | 1(9.0%) | 5(11.9%) | 0.075 |
| Pulmonary infection | 30(85.7%) | 18(90.0%) | 10(71.4%) | 12(92.3%) | 9(81.8%) | 37(88.1%) | 0.629 |
| Urinary infection | 6(17.1%) | 6(30.0%) | 2(14.3%) | 1(7.7%) | 3(27.3%) | 3(7.1%) | 0.200 |
| Abdominal infection | 5(14.3%) | 3(15%) | 0(0.0%) | 0(0.0%) | 0(0.0%) | 5(11.9%) | 0.323 |
| Cerebral vascular disease | 22(62.9%) | 12(60%) | 4(28.6%) | 7(53.8%) | 5(45.5%) | 22(52.4%) | 0.372 |
| Hypertension | 21(60.0%) | 13(65.0%) | 9(64.3%) | 8(61.5%) | 5(45.5%) | 32(76.2%) | 0.463 |
| Diabetes | 11(31.4%) | 6(30.0%) | 8(57.1%) | 7(53.8%) | 4(36.4%) | 17(40.5%) | 0.451 |
| Mechanical ventilation | 27(77.1%) | 14(70.0%) | 9(64.3%) | 10(76.9%) | 6(54.5%) | 27(64.3%) | 0.681 |
| Central venous catheter | 18(51.4%)<sup>a</sup> | 7(35.0%)<sup>a</sup> | 11(78.6%) | 11(84.6%)<sup>a</sup> | 6(54.5%) | 27(64.3%) | **0.035** |
| Foley catheter | 23(65.7%) | 16(80.0%) | 13(92.9%) | 12(92.3%) | 8(72.7%) | 32(76.2%) | 0.264 |
| Nasogastric tube | 24(68.6%) | 11(55.0%) | 12(85.7%) | 7(53.8%) | 7(63.6%) | 27(64.3%) | 0.486 |
| Continuous renal replacement therapy | 6(17.1%) | 4(20.0%) | 0(0.0%) | 2(15.4%) | 2(18.2%) | 3(7.1%) | 0.391 |
| All cause in-hospital mortality | 13(37.1%)<sup>a</sup> | 4(20.0%)<sup>a</sup> | 11(78.6%)<sup>a</sup> | 1(7.7%)<sup>a</sup> | 6(54.5%) | 13(31.0%)<sup>a</sup> | **0.001** |
| Charlson comorbidity index<sup>b</sup> | 4.72±0.81 | 3.55±0.61 | 3.91±0.63 | 4.41±0.69 | 3.88±0.46 | 2.74±0.77 | 0.232 |
| Pitt Bacteremia Score<sup>c</sup> | 3.50±0.51 | 2.80±0.64 | 2.64±0.59 | 3.92±0.54 | 3.55±0.79 | 2.81±0.73 | 0.211 |

Abbreviations: CT, complex type.
<sup>a</sup> These data were significant using pairwise analyses: CT3176 vs CT1313 (All cause in-hospital mortality; $P = 0.012$ ); CT3176 vs CT1291 (All cause in-hospital mortality; $P = 0.001$ ); CT3176 vs CT1814 (All cause in-hospital mortality; $P = 0.0003$ ); CT3176 vs The other CTs (All cause in-hospital mortality; $P = 0.004$ ); CT1814 vs CT1313 (Central venous catheter; $P = 0.049$ ); CT1814 vs CT1291 (Central venous catheter; $P = 0.011$ );
<sup>b</sup> Charlson comorbidity index. The charlson comorbidity index can range from 0 to 37. (A patient <40 who has no other medical or surgical conditions would have a score of 0; an 81-year-old with AIDS and cancer would have a score of 37).
<sup>c</sup> A Pitt bacteremia score >4 indicates severe illness. The Pitt bacteremia score is based on temperature, hypotension, mental status, and presence or absence of mechanical ventilation.
\*p values are for comparisons between CTs. Dichotomous variables are analyzed using Fisher's exact test and continuous variables are analyzed using one-way ANOVA.

### Pan-GWAS Analysis

To identify the genetic basis of CT3176, we performed a Pan-GWAS analysis between the CT3176 and non-CT3176 groups. We identified 39 genes with known functions associated with CT3176 (*P* < 0.01 adjusted with Benjamini-Hochberg’s method) (Figure 2), including virulence factors, capsule synthesis genes, antimicrobial resistance genes and multidrug efflux transporters. Among them, *ampR* had the greatest association with CT3176. However, we noted that other CTs, such as CT3178 and some CT1689 isolates (AH1689-2 and AH1689-6), also carried the *ampR* gene (Figure 1). All isolates carrying the *ampR* gene belong to KL47.

**Figure 2.**
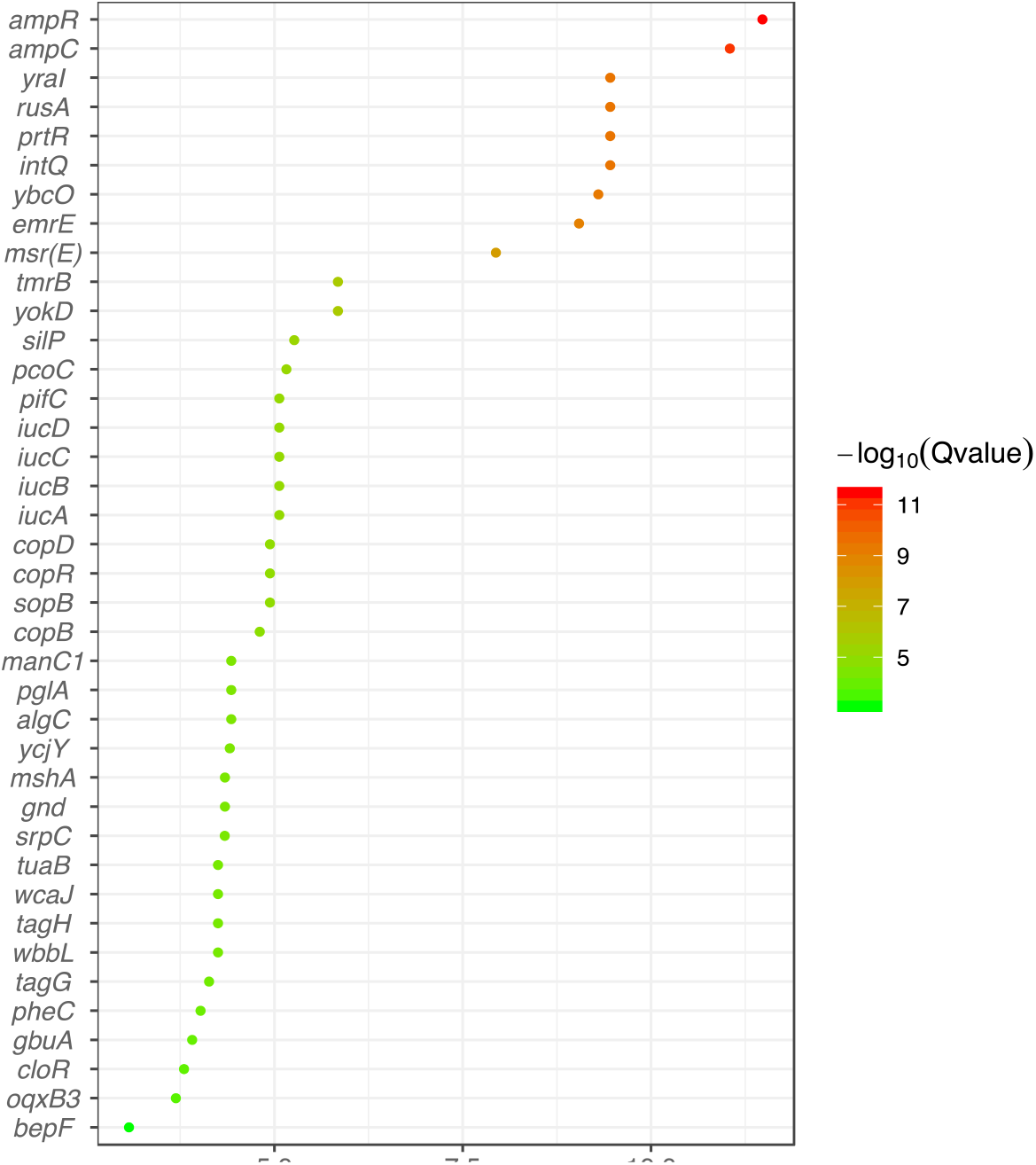
Pan-GWAS analysis between the CT3176 and non-CT3176 groups. The pangenome and associations were calculated with Roary and Scoary. A plot map was generated with ggplot2.

### Virulence in Animal Models

We aimed to identify the role of *ampR* in the virulence of non-hypermucoviscous CRKP isolates in this study. We selected three isolates (AH3176-1, AH1689-2 and AH1689-3), and tested the virulence of the isolates in a mouse model. AH3176-1 and AH1689-2 were two *ampR*^+^ isolates, while AH1689-3 was the *ampR*^−^ isolate (Figure 1). The AH1689-3 *ampR* complement mutant (AH1689-3∷*ampR*), ATCC700721 and the hypermucoviscous GM2 isolate were also included in the virulence test. GM2 belonged to ST23:KL1, which is considered a typical hvKp isolate [19]. Genomic analysis showed that *ampR* was not carried by GM2.

As shown in Figure 3A, AH3176-1 and AH1689-3∷*ampR* plus arabinose had a survival of 0% with an inoculum of 1 × 10^7^ colony-forming units (CFU) at 4 d post-infection, while 20% survival with AH1689-2, 40% survival with AH1689-3∷*ampR* without arabinose, 60% survival with AH1689-3∷pBAD33 and ATCC70721, and 80% survival with AH1698-3 was observed at 7 d post-infection. Surprisingly, the virulence of the non-hypermucoviscous *ampR*^+^ isolates AH3176-1 and AH1689-2 was comparable to that of the typical hypervirulent isolate GM2. After complementation with the *ampR* gene, survival with AH1689-3 was significantly decreased (n = 10; *P* = 0.033, log-rank Mantel-Cox test), suggesting the important role of *ampR* in virulence. Infection of mice with AH3176-1 resulted in approximately 10-100-fold higher CFU in the lungs compared to other isolates (n = 10; *P* < 0.0001, one-way ANOVA) (Figure 3B). The pathology of the lungs showed varying degrees of alveolar wall thickening, lymphocyte infiltration, and intravascular bleeding in the infection group. AH3176-1 and GM2 infection showed more pathological changes compared to ATCC700721 infection. Among them, the pulmonary pathological changes caused by AH3176-1 were the most serious, presenting as extensive pulmonary consolidation and intra-alveolar haemorrhage (Figure 3C-F). The survival of the GM2 *ampR* complement mutant was unchanged with an inoculum of 1 × 10^7^ or 1 × 10^6^ CFU (Figure 3G-H).

**Figure 3.**
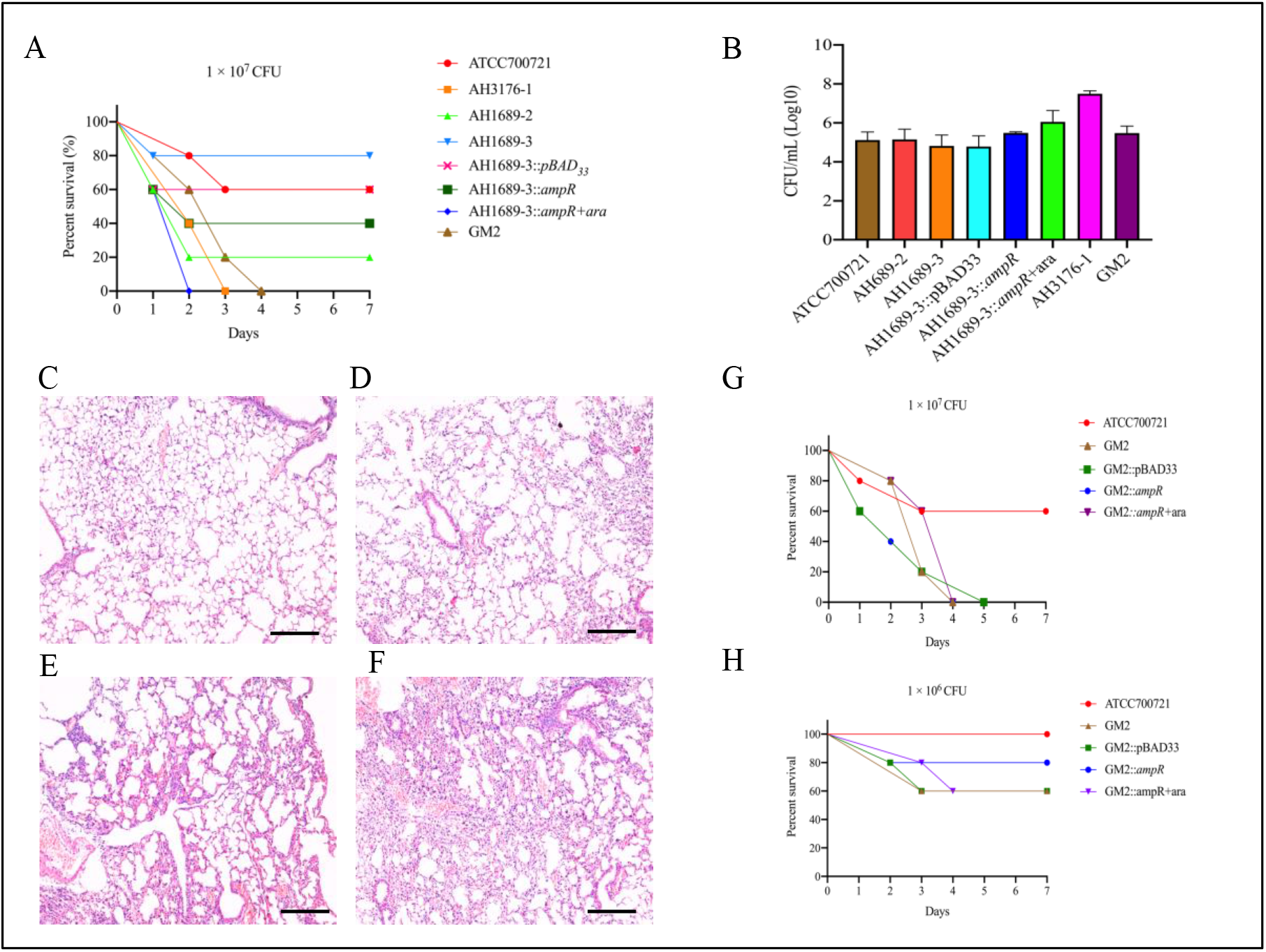
Virulence potential of *Klebsiella pneumoniae* isolates in a mouse pulmonary infection model. *A,* The effect of 1 × 10^7^ colony-forming units (CFU) of each *K. pneumoniae* isolate on survival was assessed. *B,* At 24 hours post-infection (hpi), the lungs of infected mice were harvested, and the bacterial burden was determined by CFU enumeration (n = 10; *P* < 0.0001, one-way ANOVA). Data are representative of 3 independent experiments. The lungs of mice without infection (*C*) or infected with ATCC700721 (*D*), GM2 (*E*) and AH3176-1 (*F*) were collected and stained with haematoxylin and eosin. All images are at ×40 magnification. Scale bars: 250 μM. *G,* Virulence comparison between the GM2 and GM2 *ampR* complement mutants. Abbreviation: ara, arabinose.

### Proteome Analysis, RT-qPCR and Capsule Staining

Three *ampR*^+^ (AH3176-1, AH3178-1 and AH1689-2) and three *ampR*^−^ (AH1689-3, AH1689-4 and AH1689-5) isolates were selected for proteome analysis. A total of 3093 proteins were identified in both groups based on 36727 unique peptides. Finally, we identified 24 differentially expressed proteins (DEPs). Among them, 15 were upregulated, and 9 were downregulated in the *ampR*^+^ isolates compared to the *ampR*^−^ isolates (Figure 4A-C, Supplementary Table 2). WcaJ had the highest fold change and was therefore highlighted. RT-qPCR showed that the transcription of *wcaJ* in the *ampR*^+^ isolates or *ampR* complement mutants was significantly higher than that in the *ampR*^−^ isolates (*P* = 0.0002, one-way ANOVA) (Figure 4D). The thickness of the capsule in the *ampR*^−^ isolate AH1689-3 was also increased by the *ampR* complement (Figure 4E-F).

**Figure 4.**
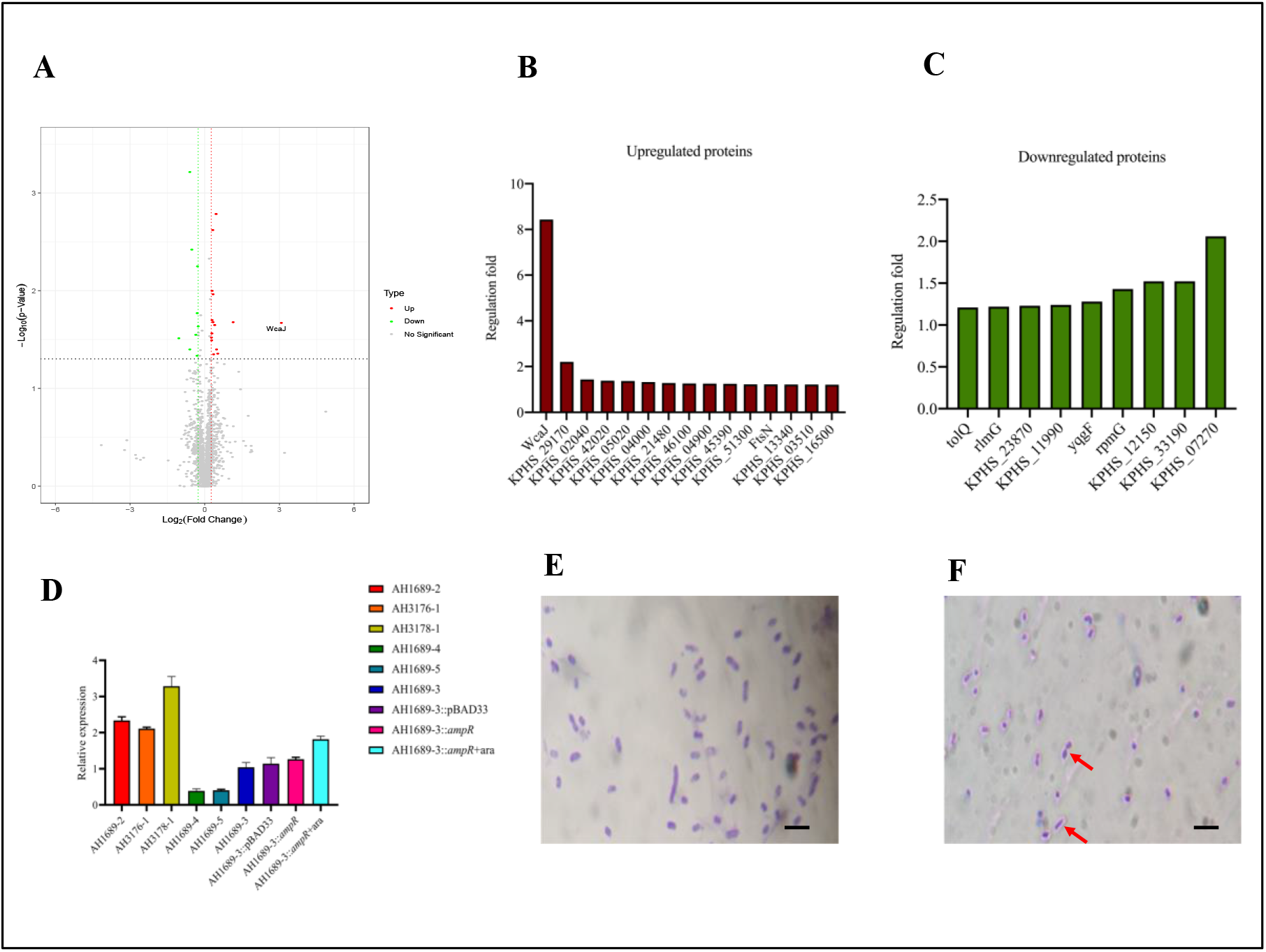
Proteome analysis, RT-qPCR and capsule staining. *A,* Volcano plot showing the comparison of quantitative protein expression between *ampR*^+^ (AH1689-2, AH3176-1 and AH3178-1) and *ampR*^−^ (AH1689-3, AH1689-4, and AH1689-5) *Klebsiella pneumoniae* isolates. Differentially expressed proteins with a fold change over 1.2 are marked in colour. *B,* Transcription level of *wcaJ* in the *ampR*^+^, *ampR*^−^ and AH1689-3 *ampR* complement isolates (*P* < 0.0001, one-way ANOVA). Capsule staining of (*C*) AH1689-3 and (*D*) AH1689-3∷*ampR* + arabinose. The red arrow points to the unstained capsule surrounding the purple-stained bacteria. Scale bars: 1 μM.

## Discussion

MLST has been the most commonly used technique for defining *K. pneumoniae* populations. With fast and affordable WGS, it is possible to compare whole genomes for isolate typing rather than just a few loci, as in traditional MLST. Genotyping with cgMLST uses thousands of alleles across the genome, resulting in a higher level of isolate discrimination. In this study, we used cgMLST to classify 135 clinical CRKP isolates and found differences between CT types based on clinical information. Due to the limited number of cases, we were unable to obtain an association between CT types and clinical outcome. However, the differences in mortality rates allowed us to further investigate the isolates belonging to CT3176. GWAS analysis showed that the *ampR* gene was significantly associated with the CT3176 isolates.

Previous studies have shown that the AmpC β-lactamase regulator AmpR, a member of the LysR family of transcription factors, also controls multiple virulence mechanisms in *Pseudomonas aeruginosa* [28, 29]. To date, only one article has reported the role of AmpR in regulating the virulence of *K. pneumoniae*. A clonal isolate of *K. pneumoniae* showed that few known virulence genes were responsible for severe infections. AmpR in these isolates was involved in the upregulation of capsule synthesis, modulated biofilm formation and type 3 fimbrial gene expression, as well as colonization of the murine gastrointestinal tract [30]. However, the virulence level of these isolates remains unclear due to the lack of data on lethal infection models. In this study, we used a pneumonia model to confirm the hypervirulence phenotype of these *ampR*-carrying isolates. Importantly, the virulence of these isolates was comparable to that of a typical hvKp isolate belonging to ST23:KL1. This would explain why patients infected with these isolates had a high mortality rate.

Previously, it was common for ST11-type *K. pneumoniae* to be resistant to carbapenems but not hypervirulent. However, in 2017, an outbreak of hospital infections caused by ST11 carbapenem-resistant hypervirulent *K. pneumoniae* isolates was reported. The hypervirulence phenotype of these isolates was due to the acquisition of an approximately 170 kbp pLVPK-like virulence plasmid by classic CRKP isolates belonging to ST11 and serotype K47 [15]. In contrast to the above report, no virulence plasmids were found in hypervirulent ST11 CRKP isolates in this study, suggesting that the hypervirulence phenotype was not due to the classical mechanism, such as RmpA and/or RmpA2 enabling bacteria to produce more capsular polysaccharides in classical hvKp [31]. Unlike the virulence plasmid, which contains multiple virulence factors, AmpR can enhance virulence independently. This was confirmed by virulence testing of the *ampR* complement strain.

WcaJ is the enzyme that initiates colanic acid synthesis and loads the first sugar (glucose-1-P) on the lipid carrier undecaprenyl phosphate. The potential role of WcaJ in the virulence of *K. pneumoniae* was defined recently [32]. The results of proteome analysis suggest that AmpR enhances the virulence of *K. pneumoniae* by regulating the initial step of capsule synthesis. As a regulator, AmpR needs to bind to the gene promoter to exert its regulatory role. To date, many capsular types have been identified in *K. pneumoniae* [33], and the absence of AmpR binding sites in some capsule types may explain why the virulence of KL1 isolates cannot be enhanced by the AmpR found in KL47 isolates.

New evidence has suggested that hypermucoviscosity and hypervirulence are different phenotypes that should not be used synonymously. Moreover, it is important to establish that a negative string test is insufficient in determining whether an isolate is hypervirulent [34]. A key finding of our work was that we identified novel non-hypermucoviscous ST11 CRKP isolates that lack most known virulence factors.

The main limitation of this study is that there are only a small number of cases; therefore, we are unable to study the relationship between the clinical outcomes and non-hypermucoviscous hypervirulent CRKP infection. Since strains with the non-hypermucoviscous phenotype are hardly distinguished in the clinic, continued surveillance and investigation of these novel strains are urgently needed.

## Supplementary Data

Supplementary materials are available at Clinical Infectious Diseases online (http://cid.oxfordjournals.org). Supplementary materials consist of data provided by the author that are published to benefit the reader. The posted materials are not copyedited. The contents of all supplementary data are the sole responsibility of the authors. Questions or messages regarding errors should be addressed to the author.

